# Cardioprotection by nicotinamide mononucleotide (NMN): Involvement of glycolysis and acidic pH

**DOI:** 10.1101/230938

**Authors:** Sergiy M. Nadtochiy, Yves T. Wang, Keith Nehrke, Joshua Munger, Paul S. Brookes

**Author notes:** Corresponding Author: Paul S. Brookes Anesthesiology and Perioperative Medicine Box 604, University of Rochester Medical Center, 601 Elmwood Avenue, Rochester, NY, 14642, USA.

## Abstract

Stimulation of the cytosolic NAD^+^ dependent deacetylase SIRT1 is cardioprotective against ischemia-reperfusion (IR) injury. NAD^+^ precursors including nicotinamide mononucleotide (NMN) are thought to induce cardioprotection via SIRT1. Herein, while NMN protected perfused hearts against IR (functional recovery: NMN 42±7% vs. vehicle 11±3%), this protection was insensitive to the SIRT1 inhibitor splitomicin (recovery 47±8%). Although NMN-induced cardioprotection was absent in *Sirt3^-/-^* hearts (recovery 9±5%), this was likely due to enhanced baseline injury in *Sirt3^-/-^* (recovery 6±2%), since similar injury levels in WT hearts also blunted the protective efficacy of NMN. Considering alternative cardiac effects of NMN, and the requirement of glycolysis for NAD^+^, we hypothesized NMN may confer protection via direct stimulation of cardiac glycolysis. In primary cardiomyocytes, NMN induced cytosolic and extracellular acidification and elevated lactate. In addition, [U-^13^C]glucose tracing in intact hearts revealed that NMN stimulated glycolytic flux. Consistent with a role for glycolysis in NMN-induced protection, hearts perfused without glucose (palmitate as fuel source), or hearts perfused with galactose (no ATP from glycolysis) exhibited no benefit from NMN (recovery 11±4% and 15±2% respectively). Acidosis during early reperfusion is known to be cardioprotective (i.e., acid post-conditioning), and we also found that NMN was cardioprotective when delivered acutely at reperfusion (recovery 39±8%). This effect of NMN was not additive with acidosis, suggesting overlapping mechanisms. We conclude that the acute cardioprotective benefits of NMN are mediated via glycolytic stimulation, with the downstream protective mechanism involving enhanced ATP synthesis during ischemia and/or enhanced acidosis during reperfusion.

## 1. Introduction

The SIRT family of NAD^+^ dependent lysine deacylases are important regulators of metabolic health [1,2]. SIRT activity is known to decline with age [3], and as such a number of NAD^+^ biosynthetic precursors are under investigation as *nutriceuticals* aimed at ameliorating diseases of aging [4,5]. Among these compounds, nicotinamide riboside (NR) and nicotinamide mononucleotide (NMN) are orally bioavailable and are currently the subject of clinical trials (NCT02812238, NCT02835664, NCT02303483, NCT02678611, etc.) [4,6].

Cardiac SIRT1 is important in several cardioprotective paradigms, including ischemic preconditioning (IPC) [7–9]. Recently, it was shown that NMN confers cardioprotection in a mouse model of ischemia-reperfusion (IR) injury, and this protection was lost in cardiac specific *Sirt1*^-/-^ mice (*CS.Sirt1*^-/-^) [10]. Although thus far the biological activity of NMN has been mostly attributed to SIRT1 stimulation [11], the role of other mechanisms including other SIRTs has received less attention. Furthermore, the NAD^+^/NADH redox couple is critically important for metabolic pathways including glycolysis and mitochondrial oxidative phosphorylation [12]. The acute effects of boosting cellular NAD^+^ levels on these metabolic pathways are poorly understood, and the role that such metabolic perturbations may have in the protective effects of NMN are unknown. Herein, we investigated alternative cardioprotective mechanisms of NMN, beyond SIRT1.

## 2. Methods

### 2.1 Animals & reagents

Chemicals and reagents were from Sigma (St. Louis MO). Wild-type C57BL6/J mice and *Sirt3*^-/-^mice [13] were bred in-house, handled according to the “NIH Guide” with food and water *ad libitum*, and used at age 8–12 weeks.

### 2.2 Langendorff perfused mouse heart IR injury model

Following tribromoethanol anesthesia (200mg/kg ip), hearts were excised and perfused in Langendorff constant flow mode (4 ml/min.) as previously described [9]. Krebs Henseleit (KH) perfusion buffer was supplemented with either (i) 5 mM glucose plus 100 μΜ palmitate-BSA, (ii) palmitate-BSA alone (glucose-free), or (iii) 5 mM galactose plus 100 μΜ palmitate-BSA [9]. Drugs (vehicle, NMN, or splitomicin (Sp)) were delivered via a port above the perfusion cannula from a 100× stock in KH for 20 min., either immediately before ischemia or at the beginning of reperfusion. Cardiac function was monitored digitally at 1 kHz (DATAQ Instruments, Akron OH) via a pressure transducer linked to a fluid-filled left ventricular balloon [9]. Following 25 or 35 min. global no-flow ischemia and 60 min. reperfusion, hearts were stained with tetrazolium chloride for infarct determination by planimetry [9]. The concentration of Sp used (10 μM) is known to block IPC and the metabolic remodeling and cytosolic lysine deacetylation that accompany it [14].

### 2.3 Cardiac metabolomics

Following 20 min. stable normoxic Langendoff perfusion accompanied by either vehicle or NMN delivery, hearts were freeze-clamped in liquid N_2_ and cardiac metabolites extracted in 80% MeOH for LC-MS/MS based metabolomics as previously described [15]. For glycolytic flux measurements, following 20 min. stable normoxic perfusion (with vehicle or NMN), glucose in KH buffer was replaced with [U-^13^C]glucose for 5 min. Hearts were then freeze-clamped in liquid N_2_ and the fractional saturation of selected cardiac metabolites with ^13^C label was determined by LC-MS/MS as previously described [15].

### 2.4 Cardiomyocytes

Primary cardiomyocytes were isolated by collagenase digestion, and intracellular pH was measured using fluorescence microscopy with the pH sensitive probe BCECF, as previously described [16,17]. A high K^+^ nigericin calibration from pH_e_ 5.6–8.4 was used to derive a Boltzmann curve fit (Origin 9, OriginLab Corp.) which was used to convert BCECF emission ratios (dual excitation 440/490 nm) to pH_i_ values. Cardiomyocyte oxygen consumption and extracellular acidification were measured in a Seahorse™ XF24 analyzer (Agilent, Santa Clara CA) either before or 10 min. after addition of NMN, as previously described [16]. Simulated IR (sIR) injury was performed as previously described, by exposing cells to 1 hr. anoxia in glucose free media at pH 6.5, followed by return to baseline conditions for 1 hr., and cell death assayed by LDH release [18,19].

### 2.5 Western blot

Vehicle or NMN treated hearts were fractionated by differential centrifugation as previously described [20], followed by separation by SDS-PAGE, transfer to nitrocellulose membranes and probing with anti-acetyl-lysine (K-Ac) or anti-SIRT3 antibodies (both from Cell Signaling Technology, Danvers MA), with enhanced chemiluminescence detection [9,13]. Purity of fractions was monitored by western blotting with antibodies against compartment-specific marker proteins.

### 2.6 Statistics

Statistical significance between groups was determined by ANOVA assuming a normal distribution followed by post-hoc Student's t-test (significance threshold p<0.05). Numbers of replicates (n) per experimental group are listed in Figure legends.

## 3. Results

### 3.1 NMN-induced cardioprotection is insensitive to SIRT1 inhibition

It was previously reported that acute pre-ischemic delivery of NMN was cardioprotective *in-vivo*, with protection lost in mice harboring cardiac-specific deletion of SIRT1 [10]. However, the *CS.Sirt1^-/-^* mice exhibited enhanced IR injury levels at baseline (~50% greater infarct vs. wild-type), which could have over-ridden the ability of NMN to protect via other mechanisms. We previously showed that cardioprotection by ischemic preconditioning (IPC) is blocked in heterozygous whole body SIRT1 knockouts (*Sirt1*^+/-^). Furthermore, both IPC itself [9] and the metabolic remodeling that accompanies it [15] can be blocked by the pharmacologic SIRT1 inhibitor splitomicin (Sp). As such, Sp is a useful tool to interrogate the role of SIRT1 in cardioprotective phenomena.

In this regard, Figures 1A and 1B show that pre-ischemic delivery of 1 mM NMN afforded significant protection against IR injury (post-IR functional recovery: NMN 42±7% vs. vehicle 11±3%; infarct size: NMN 34±4% vs. vehicle 66±4%; means ± SEM, n=6–7, p<0.01 for both parameters). However, in contrast with the *CS.Sirt1^-/-^* report mentioned above [10], we observed no effect of Sp on NMN-induced protection (p=0.71 vs. NMN alone).

**Figure 1:**
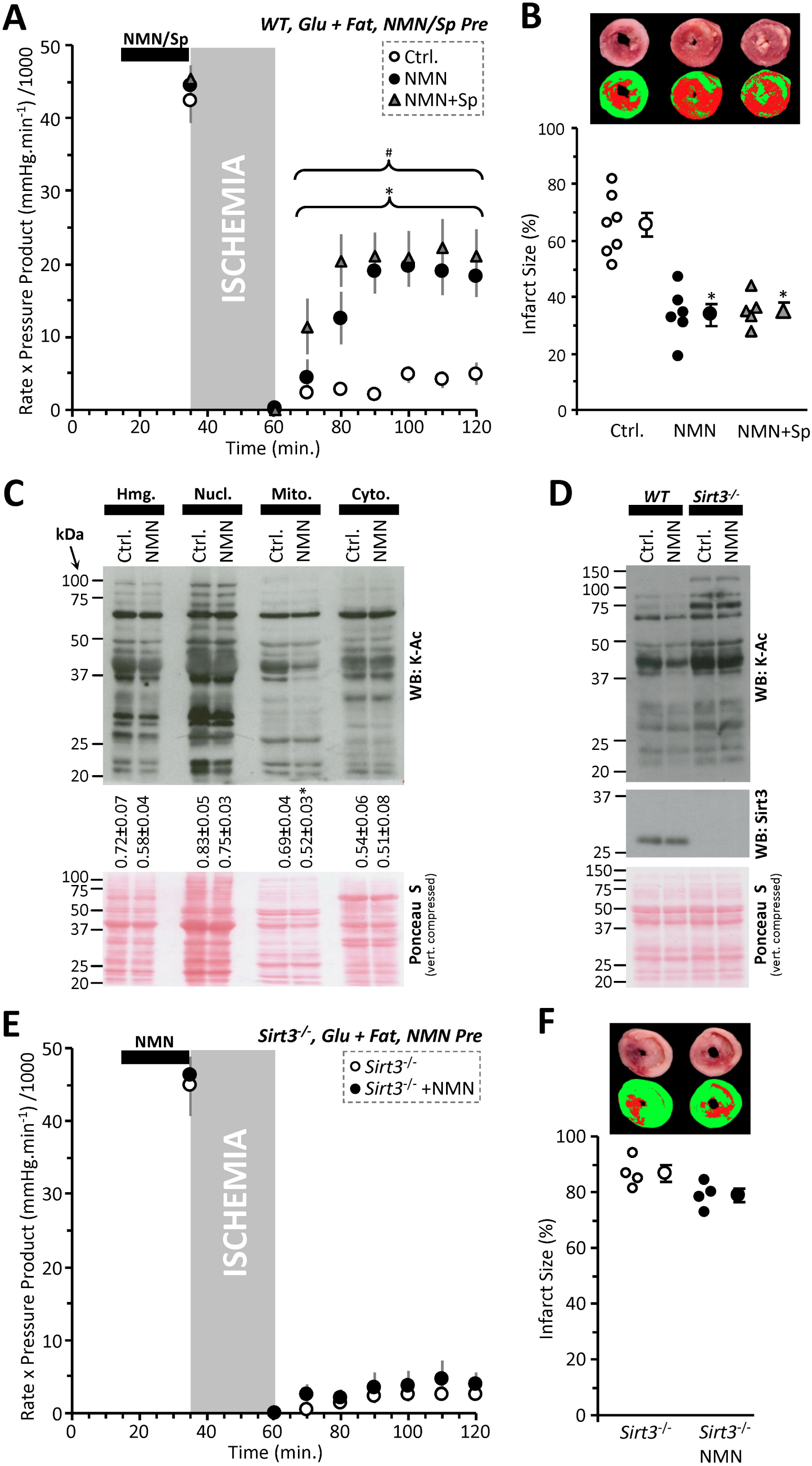
NMN Induced Acute Cardioprotection is Independent of SIRT1. **(A):** Cardiac functional data (rate x pressure product) for mouse hearts subjected to 25 min. global ischemia and 60 min. reperfusion. Prior to the onset of ischemia hearts were perfused for 20 min. with vehicle (Ctrl, white circles), 1 mM NMN (black circles), or 1 mM NMN plus 10 μM splitomicin (Sp, gray triangles). Perfusion media contained both glucose (5 mM) and fat (palmitate-BSA 100 μM) as metabolic substrates. Data shown are means ± SEM, N=5–7 animals per group, *p<0.01 for all points indicated, between Ctrl. and NMN+Sp, #p<0.01 for all points indicated between Ctrl. and NMN. **(B):** Infarction data for the hearts from panel A, determined by tetrazolium chloride staining. Images above the graph show representative infarct photographs (upper), and pseudo-colored images (lower) used for quantitation by planimetry. In graph, individual data points are shown on the left, and means ± SEM on the right, for each treatment group. *p<0.001 vs. Ctrl. **(C):** Acetyl-lysine western blot on cardiac fractions from NMN treated hearts. Hearts were perfused for 20 min. with 1 mM NMN under normoxic conditions then separated into fractions (Hmg: homogenate, Nucl: nuclear, Mito: mitochondrial, Cyto: cytosolic) by differential centrifugation. Ponceau S stained membrane (loading control) is shown below compressed vertically to save space. Numbers below the blot indicate densitometric quantitation of acetyl-lysine intensity down whole lanes, normalized to protein loading, for N=4 independent experiments (means ± SEM). Fractionation controls are shown in Figure S1A. **(D):** Acetyl-lysine western blot on mitochondrial fraction from NMN treated hearts of WT and *Sirt3^-/-^* mice. Center image shows anti-SIRT3 blot, confirming status of knockout animals. Ponceau S stained membrane (loading control) is shown below compressed vertically to save space. **(E):** Cardiac functional data (rate x pressure product) for *Sirt3*^-/-^ mouse hearts subjected to 25 min. global ischemia and 60 min. reperfusion. Prior to the onset of ischemia hearts were perfused for 20 min. with vehicle (Ctrl, white circles) or 1 mM NMN (black circles). Perfusion media contained both glucose (5 mM) and fat (palmitate-BSA 100 μM) as metabolic substrates. Data shown are means ± SEM, N=4 animals per group. **(F):** Infarction data for the hearts from panel E, determined by tetrazolium chloride staining. Images above the graph show representative infarct photographs (upper), and pseudo-colored images (lower) used for quantitation by planimetry. In graph, individual data points are shown on the left, and means ± SEM on the right, for each treatment group.

### 3.2 NMN effects on lysine acetylation

To interrogate the effects of NMN on sirtuin activity in general, we fractionated NMN-treated hearts and performed acetyl-lysine (K-Ac) western blotting. Purity of fractions is shown in Figure S1A. Figure 1C shows that NMN caused robust lysine deacetylation only in the mitochondrial fraction, not in the cytosol where cardiac SIRT1 is primarily located [21]. This result suggests that a mitochondrial sirtuin such as SIRT3 may be stimulated by NMN, and indeed Figure ID shows that NMN-induced mitochondrial deacetylation was blunted in *Sirt3^-/-^* hearts.

### 3.3. NMN cardioprotection is lost in Sirt3^-/-^ hearts, but baseline injury is greater

Consistent with a role for SIRT3 in acute NMN-induced cardioprotection, Figures IE and IF show that protection is lost in *Sirt3^-/-^* hearts. However, akin to the situation with *CS.Sirt1^-/-^* described above [10], hearts from *Sirt3^-/-^* mice also exhibited a significantly greater level of IR injury at baseline (infarct 87±3% in *Sirt3^-/-^* vs. 65±4% in WT, n=4, p<0.01). Such elevated IR injury at baseline may have precluded the ability of *Sirt3^-/-^* hearts to be protected at all. In support of this notion, extension of the ischemic time in WT hearts from 25 to 35 min. yielded a greater level of injury (9.4±2.2% functional recovery, 83.5.4±2.6% infarct, Figure S1B/C), and NMN pre-treatment failed to protect against this. As such, it is not possible to conclude that SIRT3 is required for NMN-induced cardioprotection.

### 3.4 NMN stimulates cardiac glycolysis and cytosolic acidification

We recently showed that the SIRT1 inhibitor Sp causes mild cellular alkalinization in primary mouse cardiomyocytes [22]. Thus, we hypothesized that converse stimulation of SIRT1 activity might cause acidification. As Figures 2A and 2B show, 1 mM NMN induced significant cellular acidification. In addition this same concentration of NMN was capable of protecting myocytes against simulated IR injury (Figure 2C). Notably however, the degree of cytosolic acidification seen with NMN (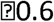 pH units) was almost double the opposing alkalinization we previously reported with Sp (+0.35 pH units) [22], suggesting additional (SIRT1-independent) mechanisms may be involved.

**Figure 2:**
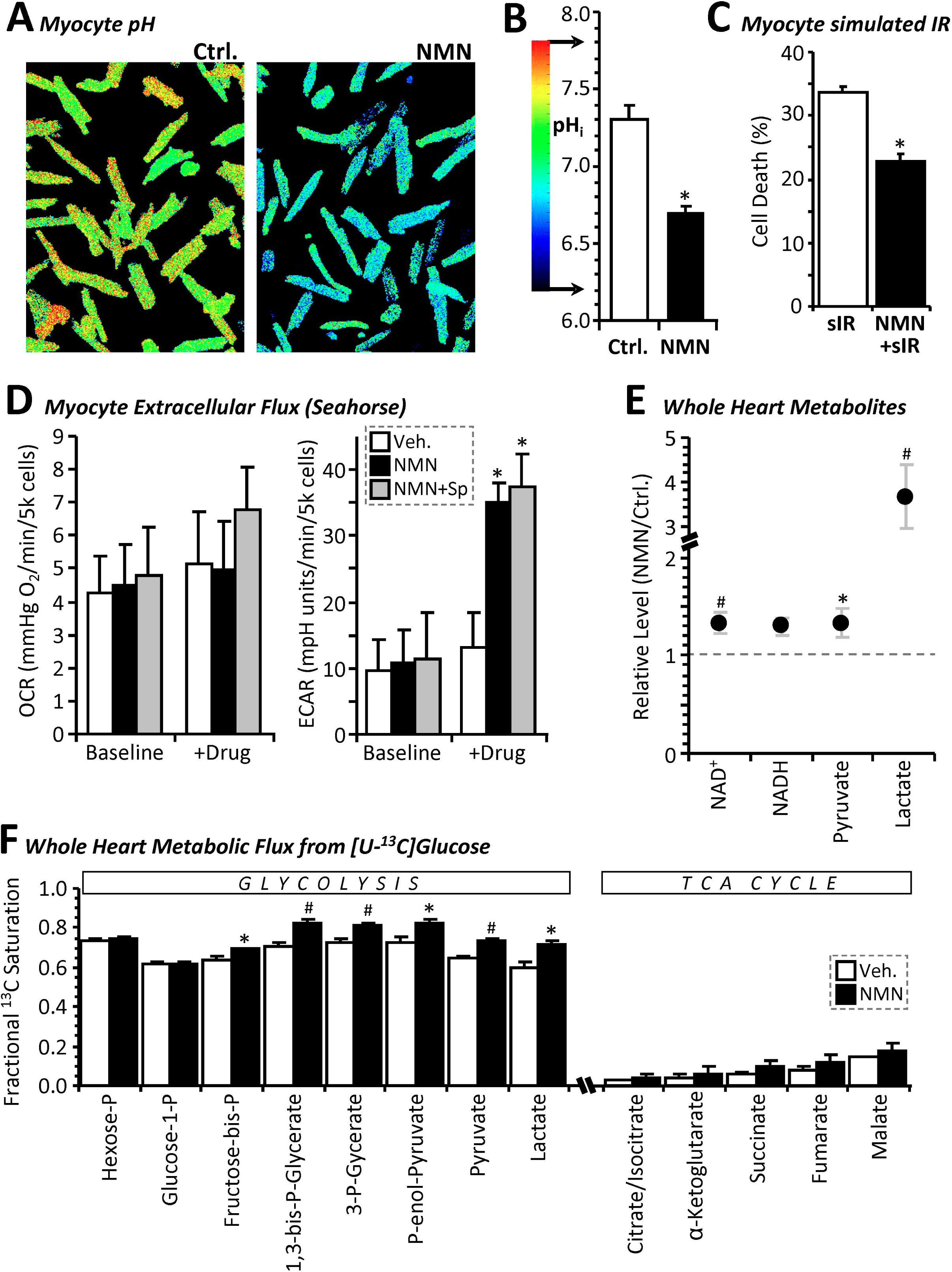
NMN Stimulates Cardiomyocyte Glycolysis Leading to Cell Acidification. **(A):** Representative fluorescent microscope images of primary adult mouse cardiomyocytes stained with the pH sensitive vital dye BCECF and treated for 40 min. with 1 mM NMN or vehicle control. Images are pseudo-colored according to the pH scale in panel B. **(B):** Graph shows average pH_i_ values from 9 independent plates of cells (means ± SEM, *p<0.001 between treatment groups). **(C):** Simulated IR injury. Cardiomyocytes were subjected to 60 min. similated ischemia followed by 60 min. simulated reperfusion (see methods), with 20 min. prior treatment with 1 mM NMN. Cell death (LDH release assay) is shown as mean ± SEM, N=5 independent cell preparations per group. **(D):** Oxygen Consumption Rate (OCR) and Extracellular Acidification Rate (ECAR) values measured via Seahorse™ XF analysis on primary adult mouse cardiomyocytes before and 10 min. after addition of vehicle (white bars) or 1 mM NMN (black bars) to the media. Where indicated, splitomicin (Sp, 10 μM) was present throughout (gray bars). Data are means ± SD for 7–8 wells per treatment group, on a single XF-24 plate. *p<0.001 between baseline and post drug treated, within a given treatment group. **(E):** Relative levels of selected metabolites in freeze-clamped perfused mouse hearts treated for 20 min. with 1 mM NMN or vehicle control. Note break in y-axis scale to accommodate lactate. Data are presented as the metabolite level in NMN treated hearts normalized to that in control hearts (means ± SEM, N=7 animals per group, *p<0.05, #p<0.01, between treatment groups). **(F):** Fractional saturation of metabolites from [U-^13^C]glucose infusion. Following vehicle control (white bars) or NMN treatment (black bars) for 20 min., glucose in KH buffer was replaced with [U-^13^C]glucose for 5 min., followed by freeze-clamp and isotopologue analysis by LC-MS/MS. Data are presented as fractional saturation – i.e., the fraction of the metabolite that is replaced by labeled metabolite within the allotted time, thus equating to metabolic flux from [U-^13^C]glucose to that point in metabolism (means ± SEM, N=5 animals per group, *p<0.05, #p<0.01 between vehicle and NMN groups).

Since NMN is a precursor for NAD^+^, and NAD^+^ is a substrate for gyceraldehyde-3-phosphate dehydrogenase (GAPDH) in glycolysis, we hypothesized that NMN-induced acidification may be partly due to direct glycolytic stimulation by mass action. Seahorse extracellular flux (XF) analysis (Figure 2D) revealed that 1 mM NMN caused a ~3-fold increase in cardiomyocyte extracellular acidification rate (ECAR) within 10 min., without significantly impacting oxygen consumption rate (OCR). Furthermore, this effect of NMN was not blocked by Sp, indicating no role for SIRT1.

In support of NMN as a direct stimulator of glycolysis, metabolomics experiments (Figure 2E) showed that delivery of 1 mM NMN to perfused mouse hearts caused a significant elevation of NAD^+^ and a similar increase in NADH (the latter suggesting NAD^+^ reduction). NMN also drastically elevated cardiac lactate, with a moderate but significant rise in pyruvate. In addition, stable isotope resolved metabolomics with [U-^13^C]glucose (Figure 2F) showed that acute NMN delivery to perfused hearts significantly elevated the fractional labeling of key glycolytic intermediates, indicative of enhanced flux. No change in TCA cycle flux was seen, although this is likely due to low fractional saturation overall, consistent with glucose not being a preferred carbon source for the heart [15]. Together, these data indicate that NMN directly stimulates glycolysis, likely by increasing the availability of its key substrate NAD^+^. The lack of effect of NMN on cytosolic lysine acetylation (Figure 1C) may indicate an inability of SIRT1 to compete for NAD^+^ with the presumably more abundant enzymes of glycolysis.

### 3.5 Requirement for glycolysis in NMN cardioprotection

Although enhanced glucose utilization is typically associated with cardiac pathology and heart failure [23–27], more recently several parallels have been drawn between cardiac glycolysis and hypoxic survival [15,28]. Thus, we hypothesized glycolytic stimulation may play a role in the cardioprotective benefits of NMN [10]. To test this, hearts were perfused without glucose (i.e., with palmitate as sole metabolic substrate). Figures 3A and 3B show that such glucose restriction blocks the protective benefit of NMN.

**Figure 3:**
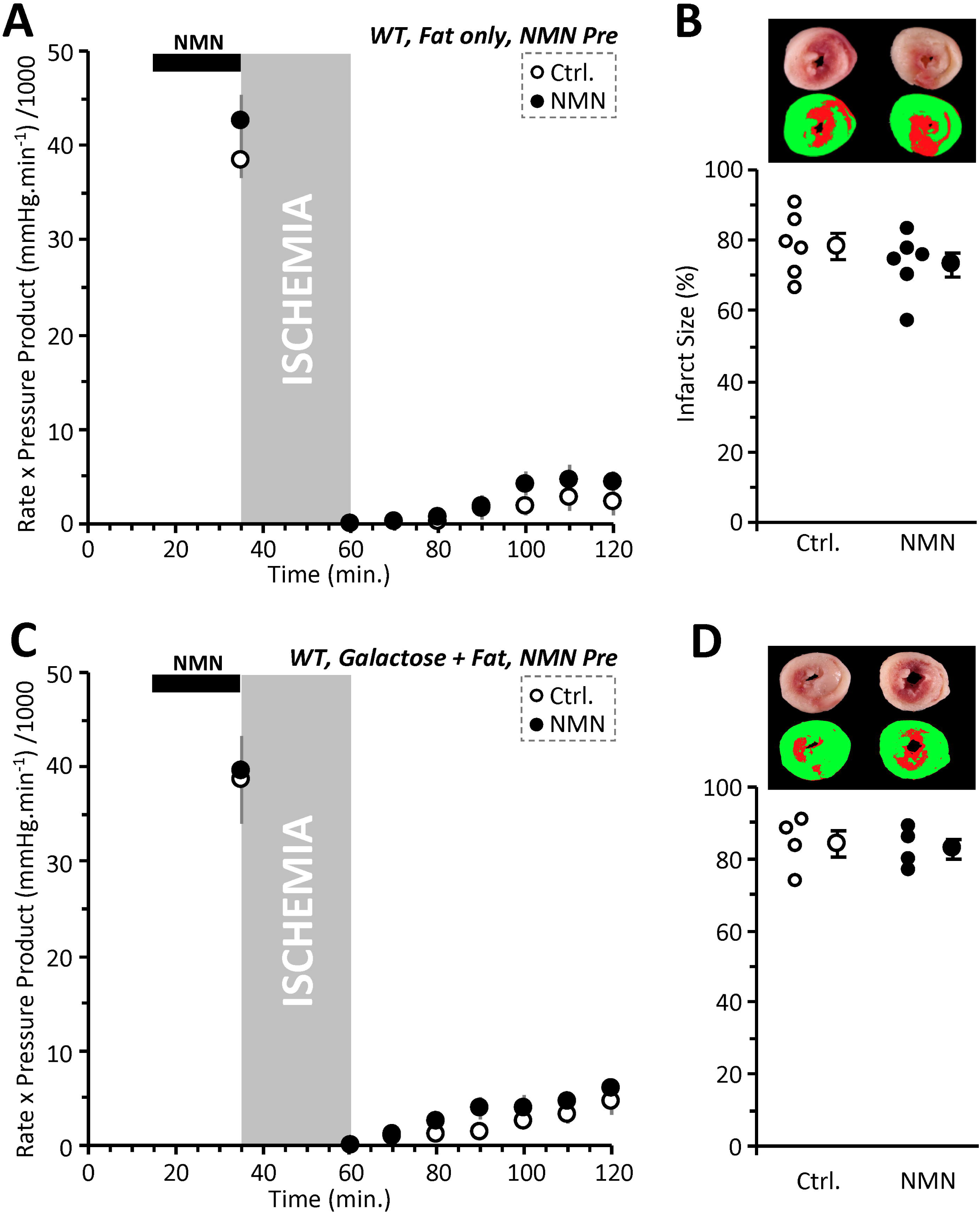
NMN Induced Acute Cardioprotection Requires Glucose. **(A):** Cardiac functional data (rate x pressure product) for mouse hearts subjected to 25 min. global ischemia and 60 min. reperfusion. Prior to the onset of ischemia (black bar at top), hearts were perfused for 20 min. with vehicle (Ctrl, white circles), or 1 mM NMN (black circles). Experiments were performed essentially as in Figure 1A, except that perfusion media contained zero glucose, such that palmitate was the sole metabolic substrate. Data shown are means ± SEM, N=6 animals per group. No significant differences were noted between Ctrl. and NMN groups. **(B):** Infarction data for the hearts from panel A, determined by tetrazolium chloride staining. Images above the graph show representative infarct photographs (upper), and pseudo-colored images (lower) used for quantitation by planimetry. In graph, individual data points are shown on the left, and means ± SEM on the right, for each treatment group. No significant differences were noted between Ctrl. and NMN groups. **(C):** Cardiac functional data (rate x pressure product) for mouse hearts subjected to 25 min. global ischemia and 60 min. reperfusion. Prior to the onset of ischemia (black bar at top), hearts were perfused for 20 min. with vehicle (Ctrl, white circles), or 1 mM NMN (black circles). Experiments were performed essentially as in Figure 1A, except that perfusion media contained 5 mM galactose instead of glucose. Palmitate-BSA was still present. Data shown are means ± SEM, N=4 animals per group. No significant differences were noted between Ctrl. and NMN groups. **(D):** Infarction data for the hearts from panel C, determined by tetrazolium chloride staining. Images above the graph show representative infarct photographs (upper), and pseudocolored images (lower) used for quantitation by planimetry. In graph, individual data points are shown on the left, and means ± SEM on the right, for each treatment group. No significant differences were noted between Ctrl. and NMN groups.

Enhancing glycolysis could conceivably elicit cardioprotection via boosting either ischemic ATP synthesis or via acidification at reperfusion. To distinguish these phenomena, we performed experiments in which hearts were perfused with galactose, which yields no net glycolytic ATP, but still permits lactate generation. In this condition, NMN pre-delivery was not protective (Figure 3C/D). These data indicate that with NMN delivered prior to ischemia, the requirement for glycolysis is one of energetics rather than enhancing acidosis during ischemia.

### 3.6 NMN-induces acid post-conditioning

A key event in IR injury is the opening of the mitochondrial permeability transition (PT) pore. During ischemia, acid pH maintains the pore in a closed state, but pH rebound upon reperfusion triggers pore opening [29–31]. As such, reperfusion with acidic media (i.e., acid postconditioning) is cardioprotective via maintaining PT pore closure into early reperfusion [32–34]. Since acidosis is a key effect of NMN (Figure 2), we hypothesized that NMN delivery at reperfusion alone may be cardioprotective. Figures 4A and 4B show that, indeed, 1 mM NMN at reperfusion was significantly protective (recovery 39±8%, infarct 28±5%, means ± SEM, n=6, p<0.01 between groups). Using the isolated primary cardiomyocyte model of simulated IR injury (see Figure 2C), we also tested the additive effects of delivering acidic buffer and/or NMN at reperfusion. These experiments (Figure 4C) revealed that while acid or NMN alone were protective, no additive (synergistic) effect was seen with both interventions. These data suggest that NMN delivery at reperfusion likely protects via acidosis. Overall, our data suggest that NMN stimulates cardiac glycolysis, and this may contribute to its cardioprotective effects, with the mechanism (energetics or acid) dependent on timing of NMN delivery relative to ischemia.

**Figure 4:**
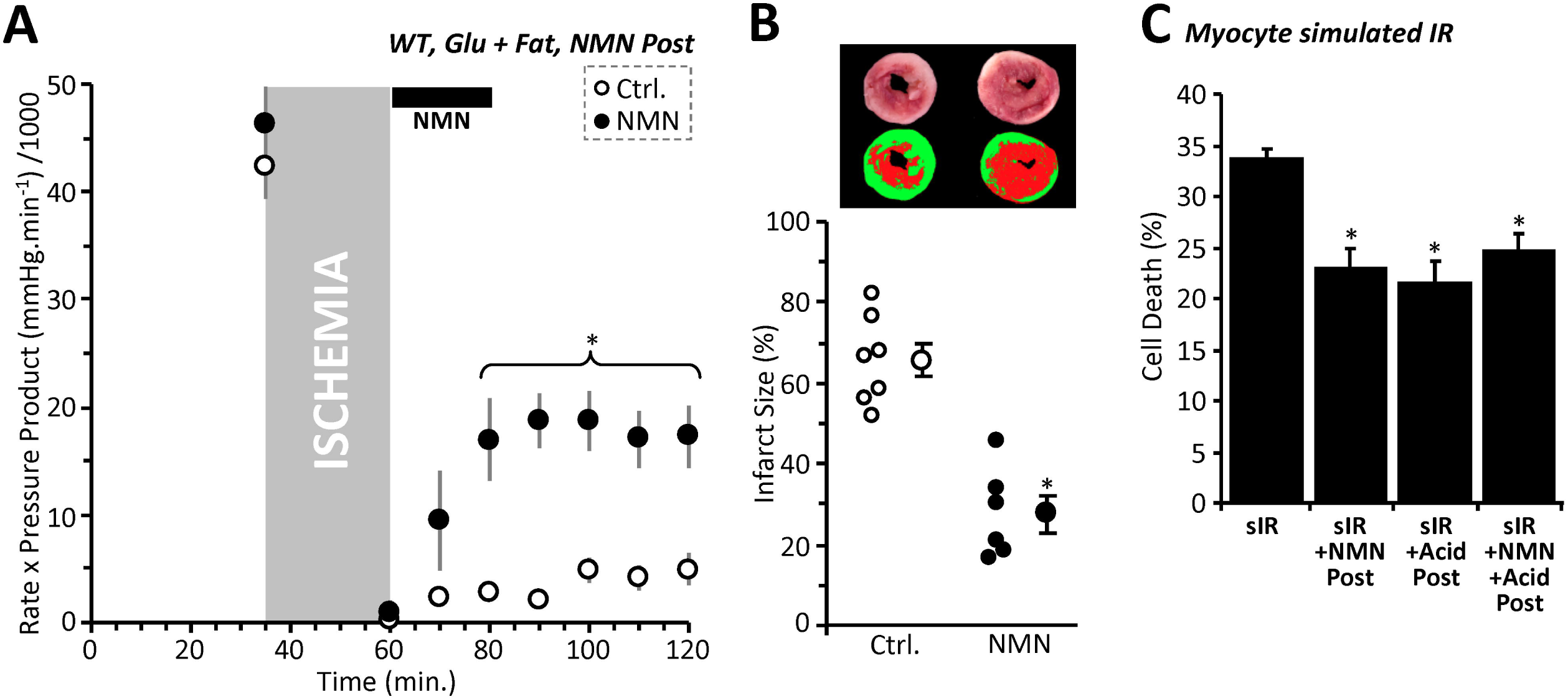
NMN is Protective at Reperfusion. **(A):** Cardiac functional data (rate x pressure product) for mouse hearts subjected to 25 min. global ischemia and 60 min. reperfusion. At the onset of reperfusion hearts were perfused for 20 min. with vehicle (Ctrl, white circles), or 1 mM NMN (black circles). Perfusion media contained both glucose (5 mM) and fat (palmitate-BSA, 100 μM) as metabolic substrates. <u>N.B.</u> Control data in this panel and panel B are the same as those in Figures 1A and B. Data shown are means ± SEM, N=6–7 animals per group. *p<0.01 for all points indicated, between Ctrl. and NMN groups. **(B):** Infarction data for the hearts from panel A, determined by tetrazolium chloride staining. Images above the graph show representative infarct photographs (upper), and pseudo-colored images (lower) used for quantitation by planimetry. In graph, individual data points are shown on the left, and means ± SEM on the right, for each treatment group. *p<0.001 between Ctrl. and NMN groups. **(C):** Simulated IR (sIR) injury. Cardiomyocytes were subjected to sIR as in Figure 2C, with delivery of either acidic media, 1 mM NMN, or both at reperfusion. Cell death (LDH release assay) is shown as mean ± SEM, N=5 independent cell preparations per group. <u>N.B.</u> sIR alone data in this panel are the same as those in Figure 2C. *p<0.01 vs. sIR alone. No significant differences noted between acid/NMN/both groups.

## 4. Discussion

The heart is conventionally viewed as a “metabolic omnivore”, but under normal conditions the bulk of cardiac ATP requirements are furnished by mitochondrial β-oxidation of fatty acids [23]. As such, a switch to favor glucose utilization is typically associated with cardiac pathology and heart failure [23–27]. However, several studies have also linked glycolysis to cardioprotection. A screen for compounds that enhance glycolysis yielded hits that were protective in models of cardiac IR injury [28]. Glycolysis is also up-regulated in IPC [15,35] and the presence of glucose is necessary for IPC [15]. Furthermore, the *pH hypothesis* of ischemic postconditioning (IPoC) posits that protection by IPoC is afforded by extending ischemic acidosis into early reperfusion, maintaining PT pore closure [32]. Together with our data showing NMN stimulates glycolysis, these findings suggest that NMN-induced cardioprotection may proceed via enhancement of glycolysis. With NMN delivery prior to ischemia, enhanced glycolysis appears to protect via enhancing bioenergetics (no protection when glycolytic ATP synthesis is nullified with galactose). On the other hand, NMN delivery at reperfusion appears to protect via acidosis (no additive effect with acidic reperfusion.

An important caveat in this study is the lack of information regarding the subcellular compartmentalization of the effects of NMN. Namely, acidification induced by NMN was cytosolic (consistent with stimulation of glycolysis), but lysine deacetylation induced by NMN was largely mitochondrial. It is not yet known whether NMN induces compartment-specific alterations in NAD^+^ or NADH pools, but the advent of novel NADH redox sensors (e.g., SoNar) may be useful in future studies aimed at determining which NAD^+^ pools are more sensitive to manipulation by compounds such as NMN [36].

While acidosis is a response of all eukaryotes to hypoxia/ischemia, the potential of acidic pH to drive protective molecular signaling events has only recently become appreciated. For example, acidic pH imparts *de-novo* activities to several metabolic enzymes, resulting in generation of unique metabolites [17,37]. In particular, the *oncometabolite* 2-hydroxyglutarate (2-HG) is generated by lactate and malate dehydrogenases under acidic conditions [17,37]. 2-HG is a competitive inhibitor of the α-ketoglutarate dependent dioxygenase family of epigenetic regulators, which includes the JmjC domain-containing histone demethylases, the TET 5-methylcytosine hydroxylases and the EGLN prolyl-hydroxylases that regulate hypoxia inducible factor (HIF) [38,39]. As such, acid pH alone can induce HIF-1α under normoxic conditions via a pathway that requires 2-HG generation [17]. The role of such a metabolic/epigenetic signaling axis in the protective effects of NMN is currently unclear.

The observation that NMN rapidly induces metabolic acidosis raises the possibility that at least some of the benefits reported for dietary NMN supplementation [4–6,12] may be attributed to such effects. A wide variety of health benefits are reported for NMN and NR supplementation, including neurological, cardiac, obesity/diabetes-related and anti-aging [12]. In many cases, such benefits may be linked to enhancing glycolysis. For example, NMN treatment has been shown to increase performance in whole body glucose tolerance tests [40], which may simply be due to enhanced glucose uptake to fuel NMN-stimulated glycolysis. Similarly, NR enhances stem cell function, with *stem-ness* being generally associated with a glycolytic metabolic state [5].

Despite the reported benefits of dietary NMN/NR supplementation, our results also urge caution regarding the widespread human use of these nutriceuticals. For example, the loss of SIRT1 activity in aging is associated with HIF stabilization and a *pseudo-hypoxic* metabolic state [41]. As such, acute acidosis resulting from NMN supplementation may activate HIF, worsening pseudo-hypoxia. Similarly, HIF activation and metabolic acidosis are well-known hallmarks of cancer [42], and indeed the same screen that identified glycolytic stimulators as protective against hypoxic injury also found numerous anti-cancer drugs as glycolytic suppressors [28]. Acid pH is also known to promote the reverse reaction of isocitrate dehydrogenase (i.e., reductive carboxylation of a-ketoglutarate to citrate), which is an important driver of lipid biosynthesis in cancer. Together with the apparent ability of NR to promote stem-ness [5], these findings suggest that NAD^+^ boosting supplements may be beneficial to cancer cells.

## 5. Conclusions

Overall, our results suggest that the nutriceutical use of NMN and NR should not overlook the classical role that NAD^+^ plays in glycolysis. GAPDH is one of the most highly and stably expressed proteins in the cell, and is often used as a “housekeeping protein” for experimental loading controls. The levels of GAPDH are likely several orders of magnitude higher than those of NAD^+^ consuming signaling enzymes such as SIRTs, PARPs, and CD38. Thus, the biological effects of large-scale acute elevations in [NAD^+^] are likely to involve glycolytic stimulation. It remains to be determined whether long-term beneficial effects of NAD^+^ precursors are mediated by repetitive transient metabolic acidosis. Although we previously reported that SIRT1 inhibition (Sp) can alkalinize cells [22], this does not appear to be sufficient to override acidification caused by NMN (Figure 2D). This effect highlights the complex nature of overlapping mechanisms that regulate cell pH, including both SIRT1 signaling and glycolytic metabolism.

## 6. Acknowledgments & conflict disclosure

This work was funded by grants from the US National Institutes of Health: RO1 HL-071158 (to PSB) and R01-HL127891 (to PSB and KWN). We thank Xenia Schafer (URMC Biochemistry) for technical support. The authors declare no financial or other conflicts of interest.

